# Discrimination between *Onchocerca volvulus* and *O. ochengi* filarial larvae in *Simulium damnosum* (*s.l.*) and their distribution throughout central Ghana using a versatile high-resolution speciation assay

**DOI:** 10.1101/046987

**Authors:** Stephen R. Doyle, Samuel Armoo, Alfons Renz, Mark J Taylor, Mike Yaw Osei-Atweneboana, Warwick N Grant

## Abstract

**Background:** Genetic surveillance of the human filarial parasite, *Onchocerca volvulus*, from onchocerciasis endemic regions will ideally focus on genotyping individual infective larval stages collected from their intermediate host, Simuliid blackflies. However, blackflies also transmit other *Onchocerca* species, including the cattle parasite *O. ochengi*, which are difficult to distinguish from the human parasite based on morphological characteristics alone. This study describes a versatile approach to discriminate between *O. volvulus* and *O. ochengi* that is demonstrated using parasite infective larvae dissected from blackflies.

**Results:** A speciation assay was designed based on genetic differentiation between *O. volvulus* and *O. ochengi* mitochondrial genome sequences that can be performed in highthroughput high-resolution melt (HRM)- or lower throughput conventional restriction fragment length polymorphism (RFLP) analyses. This assay was validated on 185 *Onchocerca* larvae dissected from blackflies captured from 14 communities in Ghana throughout 2011-13. The frequency of *O. ochengi* was approximately 67 % of all larvae analysed, which is significantly higher than previously reported in this region. Furthermore, the species distribution was not uniform throughout the study region, with 25 %, 47 % and 93 % of *O. volvulus* being found in the western-most (Black Volta, Tain and Tombe), the central (Pru) and eastern-most (Daka) river basins, respectively.

**Conclusions:** This tool provides a simple and cost-effective approach to determine the identity and distribution of two *Onchocerca* species, and will be valuable for future genetic studies that focus on parasites collected from blackflies. The results presented highlight the need to discriminate *Onchocerca* species in transmission studies, as the frequency of each species varied significantly between the communities studied.

## Background

*Onchocerca volvulus* is a human filarial parasite responsible for the disease onchocerciasis, and is transmitted between hosts by blackflies of the genus *Simulium.* Collection of blackflies throughout disease endemic regions forms a significant part of onchocerciasis surveillance campaigns to determine the prevalence and transmission of the parasite. Two approaches are commonly used to detect *O. volvulus*. The first approach, pool-screening, has been extensively used to determine the prevalence of a single parasite in a pool of low hundreds of blackfly heads [1], which can be detected using an *Onchocerca*-specific O-150 polymerase chain reaction (PCR) assay together with an *O. volvulus*-specific DNA probe. Pool-screening has been the predominant tool used for parasite surveillance by the African Programme for Onchocerciasis Control (APOC, Burkina Faso), and is suitable particularly when the prevalence of *O. volvulus* is low in areas where widespread use of ivermectin has taken place. Adaptation of the pool-screening approach to detect *O. volvulus* has recently been demonstrated using loop-mediated isothermal amplification (LAMP) targeting either the *O. volvulus* glutathione *S*-transferase 1a gene [2], or the O-150 repeat (Nankoberanyi, Doyle & Grant, personal communication; [3]), which promises to improve the throughput, cost and sensitivity of the conventional approach. Although pool-screening has the capacity to examine large numbers of flies, the primary objective of the assay is simply to determine the presence or absence of the parasite in a defined number of vectors. The second approach to detect *O. volvulus* is *via* the dissection of larvae from individual blackflies. This method pre-dates the use of pool-screening (having been in use since the initial Onchocerciasis Control Programme (OCP) in West Africa) and is still practiced widely where access to pool-screening is problematic. Dissection has the advantages that it is done at the point of fly collection and does not require the relatively complex, multi-step protocol of O-150 pool screening, and allows the developmental stage and location of the parasite(s) within the blackfly to be determined [4]. Data on larval prevalence in the vector are used together with the biting density of the fly to calculate the transmission potential of the parasite [5, 6]. It has the disadvantage, however, that it is not possible to routinely differentiate between *O. volvulus* and other *Onchocerca* species that may be transmitted concurrently by the same flies.

Genetic diagnostic tools are under development by us and others that aim to genetically define parasite populations, characterise transmission zones, and screen parasite populations for polymorphisms associated with variation in response to ivermectin [7] as a means of better defining the source and likely impact on mass drug administration effectiveness of parasite transmission in areas in which parasite transmission continues despite ivermectin distribution [8–10], i.e. genetic epidemiology. Such genetic epidemiological analyses could be integrated with, and add value to, existing entomological surveillance practices using the vector stages of the parasite as a non-invasive source of parasite material rather than microfilaria obtained from skin snips or adult parasites obtained by nodulectomy. Genetic analysis of parasites dissected from flies for the purpose of genetic epidemiology will require an efficient and sensitive molecular means to determine the species of parasite collected.

Species characterisation of dissected larvae is challenging (although not impossible) based on morphological characteristics [6, 11, 12]; it requires significant time and expertise, and is therefore relatively low-throughput. O-150-based diagnostic assays remain the predominant tool for molecular discrimination of *O. volvulus* from other *Onchocerca* species [9, 13], although sequencing of mitochondrial DNA derived amplicons has proven to be an effective approach to delineate the phylogeny of the *Onchocerca* genus [14]. However, many studies describe the prevalence of *O. volvulus* larvae in blackflies without molecular confirmation [15–18]. Given that a number of *Onchocerca* species, including the common cattle parasite *O. ochengi,* can be transmitted by the same vector [6], it is likely that *O. volvulus* prevalence may easily be overestimated, and in turn, negatively influence the predictions of disease transmission. In this study, we report the development of a versatile molecular tool to discriminate the onchocerciasis parasite *O. volvulus* from other *Onchocerca* species such as *O. ochengi* based on mitochondrial DNA (mtDNA) sequence variation. We were motivated to develop a new speciation tool, rather than using an existing O-150 assay, as a primary screening approach to discriminate *O. volvulus* from other *Onchocerca* larvae prior to high resolution melt (HRM) genotyping for population and other genetic epidemiological analyses. This approach was validated using *Onchocerca* larvae dissected from blackflies that were collected from three geographically distinct regions in Ghana.

## Methods

Blackflies were sampled from 14 communities in 5 river basins in central Ghana during 2011 (number of blackflies (*n*) = 12031), 2012 (*n* = 9706) and 2013 (*n* = 9138) using a human landing collection protocol as described previously [19]. Ethical approval was obtained for the use of human vector collectors from the Institutional Review Board of the Council for Scientific and Industrial Research, Ghana. All blackflies caught were dissected to collect *Onchocerca* larvae (129 from the head, 51 from the thorax, and 5 from the abdomen used here), after which all larvae collected were dried on microscope slides for preservation. In total, 160 infective (L3), together with 21 pre-infective L2 and 4 L1 larvae were analysed (Table 1) using the molecular techniques described, which included 14 larvae collected from 8 infected flies in the 2011 cohort, 24 larvae from 11 flies collected in 2012, and 147 larvae from 110 flies collected in the 2013 cohort. Individual larva were recovered from slides by applying 2 μl of HPLC-grade water (Sigma-Aldrich, Castle Hill, Australia) directly onto the specimen and allowing it to rehydrate, so that the larva would detatch from the slide or could be dislodged by a gentle scrape with a pipette tip. Each larva was individually transferred to 20 μl of sample lysis solution (DirectPCR Lysis reagent [MouseTail; Viagen Biotech, Los Angeles, USA] supplemented with 0.3 mg/ml (w/v) proteinase K [Roche, Mannheim, Germany]), and incubated for 18 h at 55 °C followed by 85 °C for 1 h. Larval lysates were diluted with HPLC-grade water 1/10 prior to PCR. Adult *O. volvulus* samples were obtained as previously described [10]. Adult *O. ochengi* were collected from infected cattle udders obtained from an abattoir in Ngaoundéré, Cameroon. Nodules were collected from the skin, from which worms were extracted using forceps *via* a small incision before being fixed in 95 % ethanol. DNA was extracted from individual adult *O. volvulus* and *O. ochengi* samples using an Isolate II Genomic DNA extraction kit (Bioline, Sydney, Australia) following the manufacturers’ instructions.

**Table 1.** Distribution of communities and samples used in this study

| Community | River basin | Geographical coordinates |  | Larvae processed and species identified by HRM per sample collection year<sup>a</sup> |  |  |  |  |  | Total no. of larvae per community |
| --- | --- | --- | --- | --- | --- | --- | --- | --- | --- | --- |
|  |  | Latitude | Longitude | 2011 (12,031 flies dissected) |  | 2012 (9,706 flies dissected) |  | 2013 (9,138 flies dissected) |  |  |
|  |  |  |  | *O. volvulus* | *O. ochengi* | *O. volvulus* | *O. ochengi* | *O. volvulus* | *O. ochengi* |  |
| Abua (ABA) | Pru | 7.58587 | -0.53485 | 1 | – | – | – | – | – | 1 |
| Agbelekame I (AB) | Black Volta | 8.14018 | -2.12185 | 2 | 2 | 1 | 4 | 8 | 9 L2; 27 L3 ; | 53 |
| Akanyakrom (AKA) | Black Volta | 8.289 | -2.277 | – | – | 5 | – | – | – | 5 |
| Asubende (ASU) | Pru | 8.01085 | -0.58528 | – | – | 2 | – | 5 | 2 | 9 |
| Baaya (BAY) | Pru | 8.01119 | -0.59444 | – | – | – | – | 2 | 1 L2; 12 L3 | 15 |
| Beposo (BEP) | Pru | 8.0026 | -0.57395 | – | – | – | – | 1 | – | 1 |
| Fawoman (FAW) | Pru | 8.0111 | -1.01303 | – | – | 1 | 2 | – | – | 3 |
| Fawoman-Banda (FAB) | Tombe | 8.07108 | -2.1441 | – | – | – | 6 | 2 | – | 8 |
| Kyingakrom (KYN) | Black Volta | 8.05578 | -2.03266 | – | 3 | – | – | – | – | 3 |
| Mantukwa (MAN) | Pru | 8.01064 | -1.0014 | – | – | – | – | 2 | 2 | 4 |
| New Longoro (NLG) | Black Volta | 8.08158 | -2.02096 | – | 1 | – | – | 1 | 1 L2; 8 L3 | 11 |
| Senyase (SEN) | Pru | 8.01247 | -0.59251 | – | – | – | – | 1 L1; 1 L2; 1 L3 | – | 3 |
| Tainso (TAN) | Tain | 8.0603 | -2.06538 | – | 5 | - | 3 | 1 L1; 4 L2; 7 L3 | 1 L1; 5 L2; 29 L3 | 55 |
| Wiae (WIA) | Daka | 8.19219 | -0.09429 | – | – | – | – | 1 L1; 12 L3 | 1 | 14 |
| Total larvae processed per species per year |  |  |  | 3 | 11 | 9 | 15 | 49 (3 L1; 5 L2; 41 L3) | 98 (1 L1; 16 L2; 81 L3) | 185 |

<sup>a</sup>L3 larvae unless otherwise stated

High resolution melt analysis was performed after real time PCR amplification of a 79-bp product using the Bio-Rad CFX system (Bio-Rad Laboratories, Hercules, USA). A 10 μl reaction mix was prepared by combining 5 μl of 2× SsoAdvanced Universal SYBR Green qPCR Supermix (Bio-Rad Laboratories), 0.5 μl of each of SP_OVOO_79 bp (5’-GTT TGG TTC TTG TTG ATT TG-3’) and ASP_OVOO_79 bp (5’-ACA TTA AAC CCC CTT ACC-3’) primers (0.5 μM working concentration), 2 μl of HPLC grade water, and 2 μl of the diluted larva lysate. The amplification protocol was performed as follows: 98 °C for 3 min, followed by 45 cycles of 98 °C for 10 s, 59.5 °C for 10 s and 72 °C for 10 s. The melt protocol was performed immediately after the amplification, and was as follows: 95 °C for 10 s, followed by a 65 °C annealing step from which the temperature was increased by 0.2 °C/plate read until a final temperature of 95 °C. PCR products generated from adult *O. volvulus* and *O. ochengi,* as well as a subset of 5 larvae samples from each melt curve group (representing both *O. volvulus* and *O. ochengi* larvae), were cloned into the plasmid pGem-T-Easy (Promega, Madison, USA) and sequenced to confirm the nucleotide variants (Macrogen, Seoul, Korea). All lysates, together with positive (cloned products and adult *O. volvulus* and *O. ochengi* genomic DNA samples) and no-template controls (NTC), were analysed in duplicate. Raw melt data was analysed using Precision Melt software (version 1.2; Bio-Rad Laboratories), from which the data was exported into Microsoft Excel for further quantitative analysis. Spatial (within flies i.e. head, thorax or abdomen, and between river basins), temporal (sampling year) and larval stage (L1, L2 or L3) differences between *O. volvulus* and *O. ochengi* were each examined using a 2 × 3 Chi-square (*χ*^2^) test with 2 degrees of freedom, and a *P*-value < 0.05 was considered to represent significant deviation from the null hypothesis.

A subset of post-PCR and HRM products were analysed by ApaI restriction digest to confirm the *O. volvulus-*specific digestion. A 20 μl reaction consisting of 10 μl PCR product together with 0.5 μl ApaI (50,000 U/ml; New England Biolabs, USA), 2 μl of CloneSmart buffer and 7.5 μl of HPLC-grade water was incubated for 3 h at 25 °C, after which the products were run on a 2 % agarose gel at 100 V for 50 mins and visualised using GelRed DNA stain (Biotum, Freemont, USA).

## Results and discussion

Whole mitochondrial genome alignments of the *O. volvulus* (NC_001861; [20]) and *O. ochengi* (obtained from genomic sequence available at http://www.nematodes.org/genomes/onchocerca_ochengi/; Blaxter Lab, University of Edinburgh) sequences was performed to identify a PCR compatible region that contained (i) a restriction site that was unique to one species, and (ii) additional polymorphism(s) that would result in a difference in the melting temperature between amplicons generated for each of the two species. A 79-bp region within the *nad1* gene spanning 8,655- to 8,733-bp of the *O. volvulus* mitochondrial genome was chosen that contained two synonymous and one non-synonymous nucleotide C > T transitions (C8683T [Ala > Ala, C8693T [Leu > Phe], & C8714T [Leu > Leu]) between the mtDNA alignment of the two species, one of which (C8693T; Leu > Phe) was found in an ApaI restriction site that is present in *O. volvulus* but absent in *O. ochengi* (Fig. 1a). A fourth discriminating variant was identified in this 79 bp region (G8659A; Leu > Leu); however, it was not ultimately used as it was located at the 5’ end of the optimal forward primer binding site. Melt curve analysis demonstrated a shift in Tm between the two sequences, with a mean Tm of 79.8 °C and 78.6 °C for the *O. volvulus-* and *O. ochengi*-derived sequences, respectively (Fig. 1b). HRM species determination of the unknown larval samples was performed by automated clustering of the unknown larval melt curves to curves derived from the control adult *O. volvulus* (Fig. 1b, c; green) and *O. ochengi* (Fig. 1b, c; red) samples. This translated into a significant and consistent deviation in melt curves between *O. volvulus* and *O. ochengi* samples as depicted in the fluorescence difference plots (Fig. 1c); these plots accentuate the difference in melt profiles throughout the temperature range (Fig. 1b) relative to a reference curve (set to *O. ochengi* adult control), and in turn, emphasizes differences between groups of sequences. To confirm that the difference between the two groups of melt curves were consistent with the prediction that the ApaI restriction site was present in the *O. volvulus* sequences but not in the *O. ochengi* sequences, 42 larvae-derived, 4 adult-derived and 2 cloned PCR products were analysed by restriction digest, of which a representative gel is shown in Fig. 1d. The melt curve and restriction digest data were 100 % concordant in the samples analysed, demonstrating that both approaches were equally predictive of the species in question. Finally, cloned amplicons derived from both the adult control samples and 10 larvae (5 from each melt curve group) were analysed by Sanger sequencing, confirming the discriminating variants and concordance with the reference whole mitochondrial genome sequences for the two species. This result does not, however, exclude the possibility that additional genetic variation may exist within the HRM-amplicon in either or both species of *Onchocerca* examined (or other potential *Onchocerca* species endemic to onchocerciasis regions [21–23]) that could result in melt curves that deviate from the *O. volvulus* and *O. ochengi* control sequences described here. For example, an analysis of the mitochondrial sequence of the *O. ochengi* Siisa variant [21] revealed a T,T,C haplotype that differed from the T,T,T haplotype of the adult *O. ochengi* presented (Dr Adrian Streit, personal communication). The melt temperature (Tm) of this *O. ochengi* Siisa variant is predicted to be 79.4 °C, compared to 79.7 °C for *O. volvulus* and 79.1 °C for *O. ochengi* (Tm’s were simulated using IDTDNA OligoAnalyser [https://www.idtdna.com/calc/analyzer] using the following parameters: 0.5 uM oligonucleotide, 50 mM NaCl, and 3 mM MgCl concentrations), which if present would have generated a melt curve and Tm shift intermediate to the *O. volvulus* and *O. ochengi* control sequences used. Although a single base change is unlikely to confound the interpretation between the melt profiles of the *O. volvulus* and *O. ochengi* control samples described here, this assay does offer further opportunity beyond conventional O150 assays to explore genetic variation among and within species. HRM requires that any comparison of unknown samples be made against known positive control samples; therefore, to explore potential variation beyond the assay described here, amplicons that result in melt curves that do not cluster with controls must be sequenced to identify and confirm potential novel variation present.

**Fig. 1.**
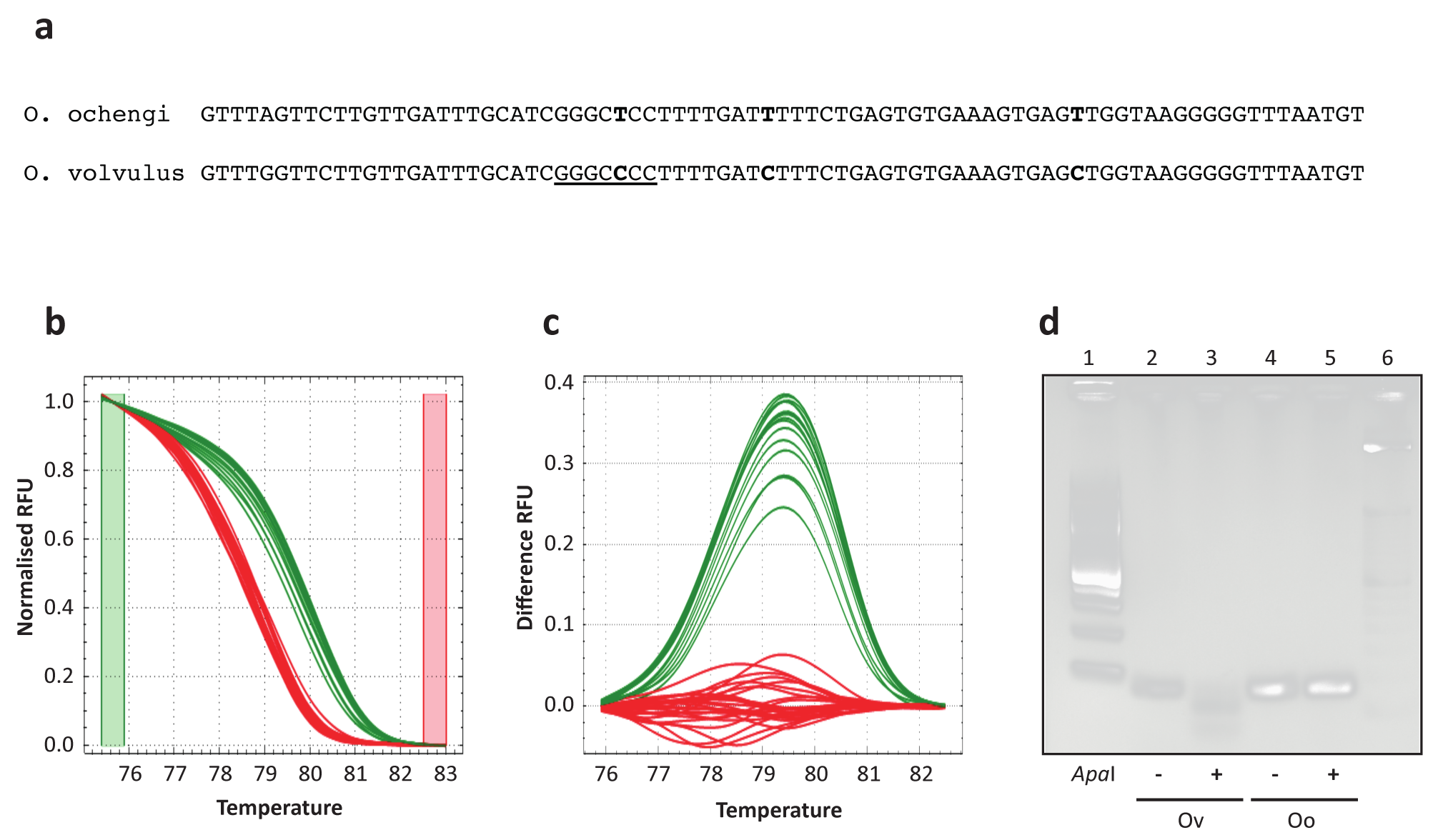
Genetic discrimination of *O. volvulus* and *O. ochengi*. **a** Sequence of the PCR 79-bp amplicon used to discriminate between *O. volvulus* and *O. ochengi*. Nucleotide differences between the two species are highlighted in bold, and the ApaI restriction site present in the *O. volvulus* sequence is underlined. **b** Normalised HRM melt curves from cloned positive control 79-bp products (pGem_Ov and pGem_Oo), adult *O. volvulus* and *O. ochengi* DNA, and 4 larvae samples from both species, each in duplicate. Curve colour of the larval samples is automatically determined based on the clustering of melt curves to the *O. volvulus* (green) and *O. ochengi* (red) adult samples. **c** The same samples are presented as in (**b**), showing HRM difference curves, which accentuate differences between the melt curve clusters in (**b**), normalised to the *O. ochengi* cluster. **d** Representative RFLP analysis of amplicons generated in the HRM assay for each species, showing digestion of the *O. volvulus* sequence, but not the *O. ochengi* sequence, with ApaI

Of the flies dissected, 166 contained *Onchocerca* larvae (Table S1); only a single larva was recovered from each of 123 flies, whereas 43 flies contained multiple larvae (Table S2; 25.90 % of total; range: 2-6). Of the flies from which larvae were processed by HRM (129 flies in total), 4 contained mixed species infections (Table S2; 14.29 % of flies with multiple larva, 3.10 % of all flies processed), i.e. they contained both *O. volvulus* and *O. ochengi* larvae, which must represent larval uptake by the blackfly from two blood meals from different hosts at different times. This is not surprising, given the prevalence of cattle and human cohabitation, and that blackflies will seek a blood meal from either host [24, 25]. The overall proportion of each larval stage was similar between both species (Table S3; *χ*^2^ = 3.397, *df* = 2, *P* = 0.091), however, the spatial distribution of each stage within the fly was not equal; although a greater proportion of L3 were found in the head relative to the thorax in both species, the difference between both head and thorax was greatest in *O. volvulus* with a higher proportion of L3 collected from the head (Figure S1, Table S3; *χ*^2^ = 6.822, *df* = 2, *P* = 0.016). This finding suggests that while the rate of development within the fly may be similar between the species, *O. volvulus* may migrate towards the head of the fly earlier that *O. ochengi*. These observations (co-infection and likely equivalent transmission potential [based on prevalence of L3]), do suggest that the human and bovine hosts are constantly exposed to both parasites which, in turn, raises an interesting question in regard to the number of times host-switching may have occurred between human and cattle (a speciation hypothesis whereby the most recent common ancestor of *O. ochengi* and *O. volvulus* was a cattle parasite that established in the human host [26]).

The distribution of both species throughout the sampling region was not uniform (Fig. 2). Although no temporal differences between sampling years for each species was seen (Table S2: *χ*^2^ = 1.075, *df* = 2, *P* = 0.292), a significant spatial trend was observed that suggested that the western communities sampled had a lower prevalence of *O. volvulus* (22.96 %, *n* = 135; Black Volta, Tain and Tombe river basins), a roughly equal prevalence of *O. volvulus* and *O. ochengi* in the central Pru river basin (47.22 % *O. volvulus*; *n* = 36), and a higher prevalence of *O. volvulus* (92.86 %, *n* = 14) in the eastern-most Daka river basin (Table S4; *χ*^2^ = 32.145, *df* = 2, *P* = 5.234e^−8^). These results are not necessarily in contrast to a recent investigation of persistent *O. volvulus* with negligible *O. ochengi* transmission in Ghana [9]. Much of that study was focused on two southern Ghanaian communities, and very few or no larvae were found in the two communities that were shared with this study (Asubende and Agborlekame, respectively); moreover, 3 of the 7 communities used by Lamberton et al. were east of the Black Volta Lake, which would be consistent with high *O. volvulus* prevalence observed in the eastern-most community (Wiae) sampled here. We speculate that the high infection rate reported and difference in prevalence of *O. ochengi* in a number of study regions presented here is correlated with the high numbers of cattle in the north-western river basins, particularly during the dry season (December to April). However, given that the sample size of larvae for many communities was low (median = 6.5, range = 1–55 larvae/community; limited by the number of larvae present in the blackfly populations and therefore by the number of blackflies screened), further sampling is required to support these findings, particularly in the Daka river basin where *Onchocerca* larvae were obtained from only a single community out of the five communities in which flies were sampled.

**Fig. 2.**
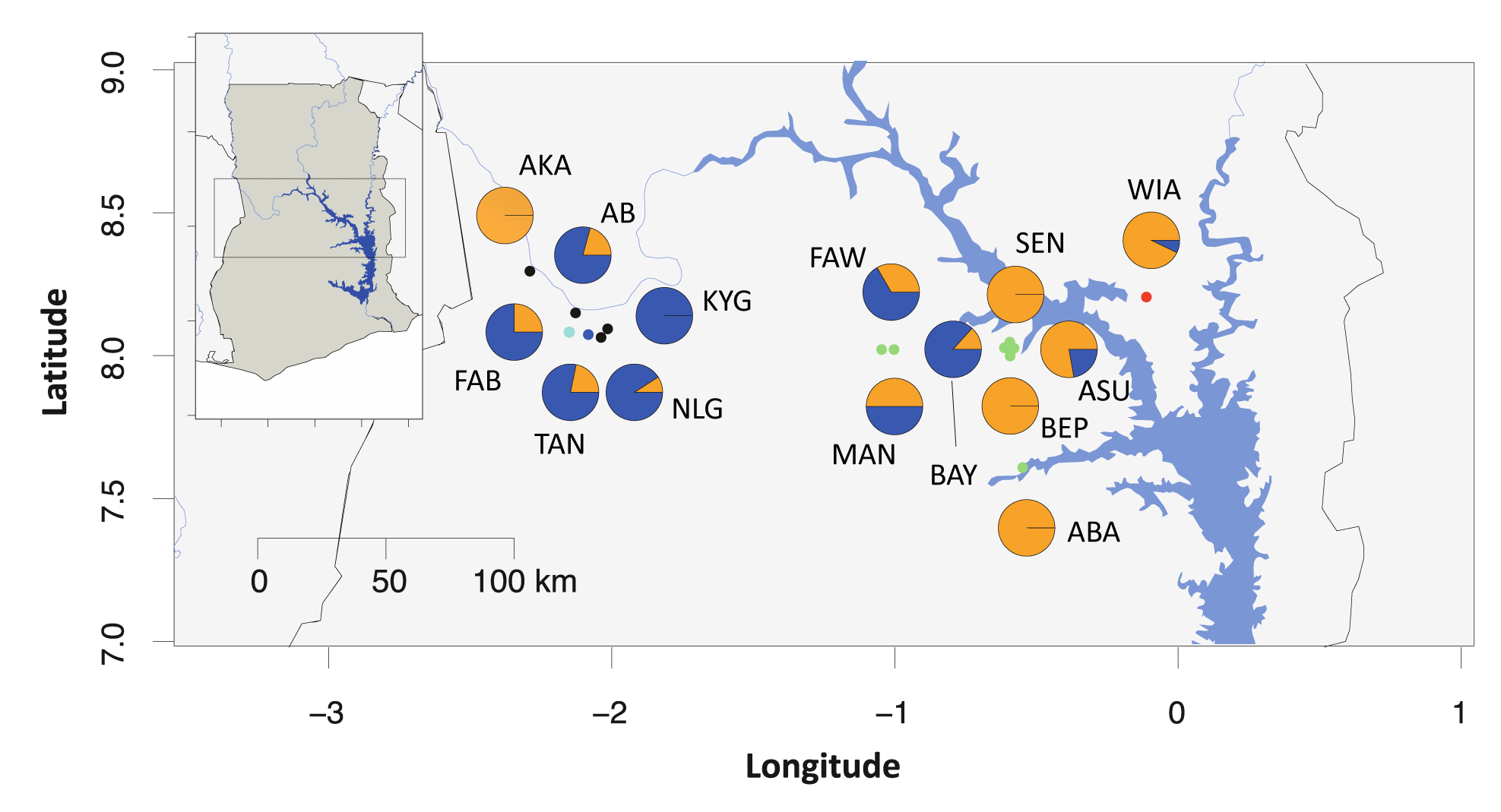
Distribution of *O. volvulus* and *O. ochengi* throughout Ghana. Pie charts depict relative prevalence of *O. volvulus* (orange) and *O. ochengi* (purple) in each community. Communities sampled in central Ghana (see insert) and the number of samples analysed are as follows: Abua (ABA; *n* = 1), Agbelekame I (AB; *n* = 54), Akanyakrom (AKA; *n* = 5), Asubende (ASU; *n* = 9), Baaya (BAY; *n* = 15), Beposo (BEP, *n* = 1), Fawoman (FAW; *n* = 3), Fawoman-Banda (FAB, *n* = 8), Kyingakrom (KYN; *n* = 3), Mantukwa (MAN; *n* = 4), New Longoro (NLG; *n* = 11), Senyase (SEN; *n* = 3), Tainso (TAN; *n* = 55), Wiae (WIA; *n* = 14). The colour coding of communities reflect the river basin in which they lie; Black Volta (black), Tain (dark blue), Tombe (light blue) Pru (green) and Daka (red)

## Conclusions

A simple but versatile tool that discriminates between *O. volvulus* and *O. ochengi* single larvae obtained by dissection of the blackfly intermediate host is described. A HRM approach offers a number of advantages over traditional O-150-based assays for the speciation of larvae collected by dissection, including (i) the ability to detect and explore novel genetic variation, either within or between Onchocerca species, and (ii) a rapid, single tube diagnostic assay format that can be run individually or in plate format (qPCR platform dependent). Note that this assay does not compete with the pool-screening O150 assay designed to detect single larvae in pools of blackflies but is designed as a component of a suite of genetic epidemiology tools under development in our laboratory to include “population of origin” and “likely ivermectin response phenotype” assays. By exploiting the multicopy nature of the mitochondrial genome (in the same way that O150 assays target a highly abundant repetitive element in the genome), this assay is very sensitive, requiring only 1/100^th^ of a single larva, thus leaving sufficient material for further genetic epidemiological analysis. Although *O. volvulus* and *O. ochengi* samples could be distinguished easily *via* HRM or RFLP assays, no effort was made to identify other described *Onchocerca* species that may coexist in onchocerciasis endemic regions [21–23]; further work is required to characterise melt profiles and/or restriction sites from additional *Onchocerca spp.* voucher specimens or sequences as they become available. In the context of onchocerciasis entomological surveillance by dissection, which aims to detect transmission of the human pathogen, and the genetic epidemiological tools under development, this study demonstrates the need to accurately and efficiently determine the identity of larvae in the vector in order to correctly estimate parasite transmission. In Ghana, nationwide sampling of blackfly populations is necessary to determine the distribution of *Onchocerca* spps. Implementation of genetic tools such as those described here will inform the appropriate control strategy needed by onchocerciais control programmes as they strive towards onchocerciasis elimination in Ghana.

## Additional files

**Additional file 1. Figure S1.** Proportion of *O. volvulus* (orange) and *O. ochengi* (purple) larval life stages (L3, L2, & L1) found during dissection of the blackfly head, thorax and abdomen. Values reported are a percentage of the total larvae found within each species.

**Additional file 2. Table S1.** Summary of microscope slides containing larvae dissected from blackflies, including location, date, life stage of larvae and location within the fly from which the larvae were dissected. **Table S2**. Summary data and analysis of larvae recovered and processed per blackfly, by species, community and geographic location. **Table S3**. Summary data and analysis of larvae collected per blackfly per year. **Table S4**. Summary data and analysis of larvae dissection from blackflies, by species, life stage (L3, L2, or L1), and tissue (head, thorax, or abdomen).

**Abbreviations**
APOC: African Programme for Onchocerciasis Control
HRM: high-resolution melt
LAMP: loop-mediated isothermal amplification
mtDNA: mitochondrial DNA
OCP: Onchocerciasis Control Programme
PCR: polymerase chain reaction
RFLP: Restriction fragment length polymorphism
Tm: melt temperature

### Acknowledgments

We thank Mr. Francis Veriegh for contributing to the blackfly dissection, and PD Dr Adrian Streit, Dr Annette Kuesel and Dr Aime Adjami for helpful suggestions toward the manuscript. We also thank Dr Mark Blaxter and Dr Benjamin Makepeace for access to the unpublished *O. ochengi* genome sequence, which was generated with support of the EU-funded programme “Enhancing Protective Immunity Against Filariasis”.

### Ethics approval and consent to participate

Ethical approval was obtained for the use of human vector collectors from the Institutional Review Board of the Council for Scientific and Industrial Research, Ghana.

### Consent for publication

Not applicable.

### Availability of data and material

All data analysed is presented in figures and tables within the manuscript and its additional files.

### Funding

This investigation received financial support from TDR, the Special Programme for Research and Training in Tropical Diseases, co-sponsored by UNICEF, UNDP, The World Bank and WHO, the German Research Foundation (DFG Re 1536/5-1) and the European Foundation Initiative for Neglected Tropical Diseases. The funding bodies did not play any role in the design of the study, collection, analysis, interpretation of data or in writing of the manuscript.

### Author’s contributions

SRD designed the study, performed the molecular biology and data analysis, and drafted the manuscript. MYO-A, MT and SA designed the entomological study for transmission assessment, SA was involved in the collection and dissection of blackflies, and preparation of larvae samples. MYO-A and MT was involved in the coordination of sample collection. AR provided samples. WG participated in the analysis and helped draft the manuscript. All authors read and approved the final manuscript.

### Author details

^1^Department of Animal, Plant and Soil Sciences, La Trobe University, Bundoora, 3086, Australia

^2^Council for Scientific and industrial Research - Water Research Institute, Accra, Ghana

^3^Institute of Evolution and Ecology, Department of Comparative Zoology, University of Tübingen, Auf der Morgenstelle 28, 74074, Tübingen, Germany

^4^Department of Parasitology, Liverpool School of Tropical Medicine, Liverpool, United Kingdom

^5^Wellcome Trust Sanger Institute, Wellcome Genome Campus, Hinxton, Cambridgeshire, CB10 1SA, United Kingdom

